# Chemotherapy induces tissue NAD^+^ loss, and downregulation of NAD^+^ biosynthetic enzyme *Nrk2* marks muscle wasting

**DOI:** 10.64898/2026.07.11.736679

**Authors:** Noora Pöllänen, Caleb J. Gammon, Fabrizio Pin, Joshua R. Huot, Roberta Sartori, Fabio Penna, Juha Hulmi, Andrea Bonetto, Eija Pirinen

**Author notes:** **Corresponding author:** Eija Pirinen.

## Abstract

**Background:** Aberrant NAD^+^ metabolism has been implicated in the pathogenesis of cancer cachexia, highlighting this pathway as a potential therapeutic target to mitigate skeletal muscle wasting. However, it remains unclear whether chemotherapeutic agents contribute to the onset of cachexia by disrupting NAD^+^ metabolism. Here, we investigated the effects of commonly used chemotherapy regimens on NAD^+^ metabolism in skeletal muscle and liver of healthy mice.

**Methods:** Healthy mice were treated with either 2-week regimens of folfiri or cisplatin, or 5-week regimens of folfiri or folfox, with vehicle-treated mice serving as controls. Cachexia-related outcomes were assessed, while skeletal muscle and liver tissues were analyzed for NAD metabolites and markers of NAD^+^ metabolism. Given the consistent downregulation of the NAD^+^ biosynthetic enzyme *Nrk2* in cachectic chemotherapy-treated mice, we examined skeletal muscle *Nrk2*/*NRK2* expression across published murine and human cachexia datasets, and in additional models of muscle wasting and hypertrophy.

**Results:** NAD^+^ loss was observed in atrophic muscle following administration of cisplatin (2-week treatment; −14% vs controls, p=0.047) and folfiri (5-week treatment; −18%, p=0.069). In contrast, muscle NAD^+^ levels were preserved in non-atrophic groups (2-week folfiri and 5-week folfox). Muscle *Nrk2* was the most responsive NAD^+^ biosynthetic enzyme, showing consistent downregulation across chemotherapy models with ongoing or developing muscle loss: cisplatin (−93%, p<0.001), folfiri (−84%, p<0.001) and folfox (−92%, p<0.001). In the liver, NAD^+^ levels declined after prolonged 5-week folfiri (−20%, p=0.013) and folfox (−15%, p=0.043) treatments. These changes were accompanied by distinct alterations in NAD^+^ biosynthesis pathways, indicating treatment-specific reorganization of hepatic NAD^+^ metabolism. Cross-study analyses revealed early and consistent skeletal muscle *Nrk2* downregulation across multiple murine cachexia models and human inactivity studies, whereas cachexia-targeted interventions in rodents and resistance training in humans increased its expression.

**Conclusions:** These findings demonstrate that chemotherapy distrupts tissue NAD^+^ metabolism, with skeletal muscle NAD^+^ loss accompanying muscle atrophy and hepatic NAD^+^ levels declining after prolonged treatment. The early and robust responsiveness of muscle *Nrk2* expression to changes in muscle mass underscores its potential as a dynamic indicator for predicting treatment-induced changes in muscle mass. Together, these results provide new molecular insight into the metabolic basis of chemotherapy-induced muscle wasting and support further investigation of NAD^+^-targeted strategies in this context.

## Introduction

Chemotherapy is a cornerstone in the treatment of various cancers and, while frequently adopted for tumor control, it is often accompanied by adverse effects (1). Beyond common side effects, such as nausea, fatigue and hair loss, chemotherapy has been shown to exacerbate cachexia, a debilitating condition of involuntary skeletal muscle and body weight loss (2). As ∼50-80% of cancer patients develop cachexia (3) characterized by weakness, fatigue, reduced quality of life and increased mortality, susceptibility to the toxicities of anticancer therapies may be further amplified. This creates a vicious cycle that necessitates balancing treatment efficacy with the management of progressive wasting. These challenges underscore the need for integrated therapeutic strategies that address both tumor control and the metabolic decline that weakens patients during treatment.

To tackle these challenges, it is crucial to elucidate the pathophysiological mechanisms and therapeutic strategies for cancer- and chemotherapy-induced cachexia. The molecular alterations induced by chemotherapy in skeletal muscle remain incompletely identified and appear to differ, at least in part, from those induced by the tumor itself (4,5). Cancer triggers catabolic and inflammatory signaling, as well as metabolic reprogramming that drive muscle wasting. Chemotherapy further disrupts muscle homeostasis by activating pro-atrophic signaling pathways, suppressing anabolic processes, and inducing mitochondrial dysfunction and metabolic derangements, ultimately accelerating muscle atrophy (4–6). Hence, understanding the shared and distinct tumor- and chemotherapy-specific mechanisms is essential for developing targeted interventions to prevent or mitigate cachexia throughout cancer treatment.

Our previous work has identified a deficiency of nicotinamide adenine dinucleotide (NAD^+^), a critical coenzyme for multiple cellular process, including energy metabolism and DNA repair, in skeletal muscle and liver as a potential pathophysiological mechanism underlying cancer cachexia. Such deficiency was observed across several murine models of cancer cachexia (7,8). Importantly, our *in vivo* studies demonstrated that NAD^+^ repletion exerted a therapeutic effect against both cancer- and cancer plus chemotherapy-induced cachexia (8). Additionally, we have identified the downregulation of NAD^+^ biosynthetic enzyme nicotinamide riboside kinase 2 (*Nrk2*, also known as *Nmrk2*, *Itgb1bp3* or *Mibp*) in skeletal muscle as a common feature of cancer- and chemotherapy-induced cachexia in mice and humans (7,8). Nrk2 is a muscle-specific enzyme that is normally activated upon stress to support NAD^+^ biosynthesis (9), and it is involved in other processes commonly disrupted in cachexia, including extracellular matrix remodeling, myogenesis and muscle regeneration (10). Paradoxically, despite these protective roles, muscle *Nrk2* is suppressed in cachexia, potentially compromising NAD^+^ homeostasis and other physiological functions and thereby contributing to muscle wasting. Collectively, these findings suggest that dysregulated NAD^+^ metabolism represents a mechanistically relevant and therapeutically actionable pathway for counteracting muscle wasting in cancer- and chemotherapy-induced cachexia.

Currently, our understanding of how chemotherapeutic agents specifically modulate NAD^+^ metabolism remains unclear. Outside skeletal muscle tissue, previous studies have reported cisplatin-induced NAD^+^ loss in the small intestine (11), and systemic NAD^+^ depletion upon doxorubicin administration (12). Notably, *Nrk2* emerged as the most downregulated transcript in skeletal muscle already following a single dose of doxorubicin (13). Altogether, these observations suggest that chemotherapy modulates NAD^+^ metabolism and results in *Nrk2* downregulation, that may contribute to tissue-specific toxicity and muscle wasting. This highlights the need to further elucidate the effects of chemotherapeutic agents on NAD^+^ metabolism and *Nrk2* expression during muscle atrophy.

In this study, we first explored the impact of cisplatin, folfox, and short- and long-term folfiri chemotherapies on tissue NAD^+^ metabolism in healthy mice. Our findings reveal chemotherapy-induced NAD^+^ loss and downregulation of *Nrk2* in skeletal muscle, recapitulating metabolic alterations previously observed in cancer cachexia. Building on these observations, we conducted a more comprehensive analysis of skeletal muscle *Nrk2* expression across diverse muscle-wasting conditions. This approach suggest that skeletal muscle *Nrk2* expression may serve as a sensitive indicator of muscle mass changes and holds promise as a predictive marker of cachexia-associated muscle wasting.

## Methods

### Animal experiments

All animal experiments were approved by the Institutional Animal Care and Use Committee at the Indiana University School of Medicine. The animals were cared for in compliance with the Declaration of Helsinki and the National Institutes of Health Guidelines for Use and Care of Laboratory Animals. The animals were maintained on a regular dark-light cycle of 12:12 h with controlled temperature (20–23 °C), humidity (40–60%) and had *ad libitum* access to food (2018 Teklad global 18% protein rodent chow) and water during the whole experimental period.

Four chemotherapy-based *in vivo* experiments were conducted with 8-week old CD2F1 wild-type mice (Inotiv, West Lafayette, IN) (Fig. 1A).

1. male mice (N=5) received a single injection with folfiri chemotherapy (50mg/kg 5-fluorouracil, 90mg/kg leucovorin, 24mg/kg irinotecan), administered intraperitoneally (i.p.). Control mice (N=5) received isovolumetric amounts of vehicle, as previously described (Barreto et al., 2016). Animals were sacrificed 24 hours later.
2. male mice (N=5) received folfiri chemotherapy (same doses as above, i.p.), twice weekly for 2 weeks. Control mice (N=5) received same amounts of vehicle, as in our previous publication (5).
3. female mice (N=5) were administered either folfiri (same doses as above, i.p., twice weekly) or folfox (50mg/kg 5-fluorouracil, 90mg/kg leucovorin, 6mg/kg oxaliplatin, i.p., once weekly). Control mice (N=5) received the vehicle only.
4. male mice (N=8) were treated with cisplatin (2.5 mg/kg, i.p.) daily for 1 week, followed by administration every other day for an additional week. Control mice (N=5) received equivalent volumes of vehicle (14).

**Figure 1.**
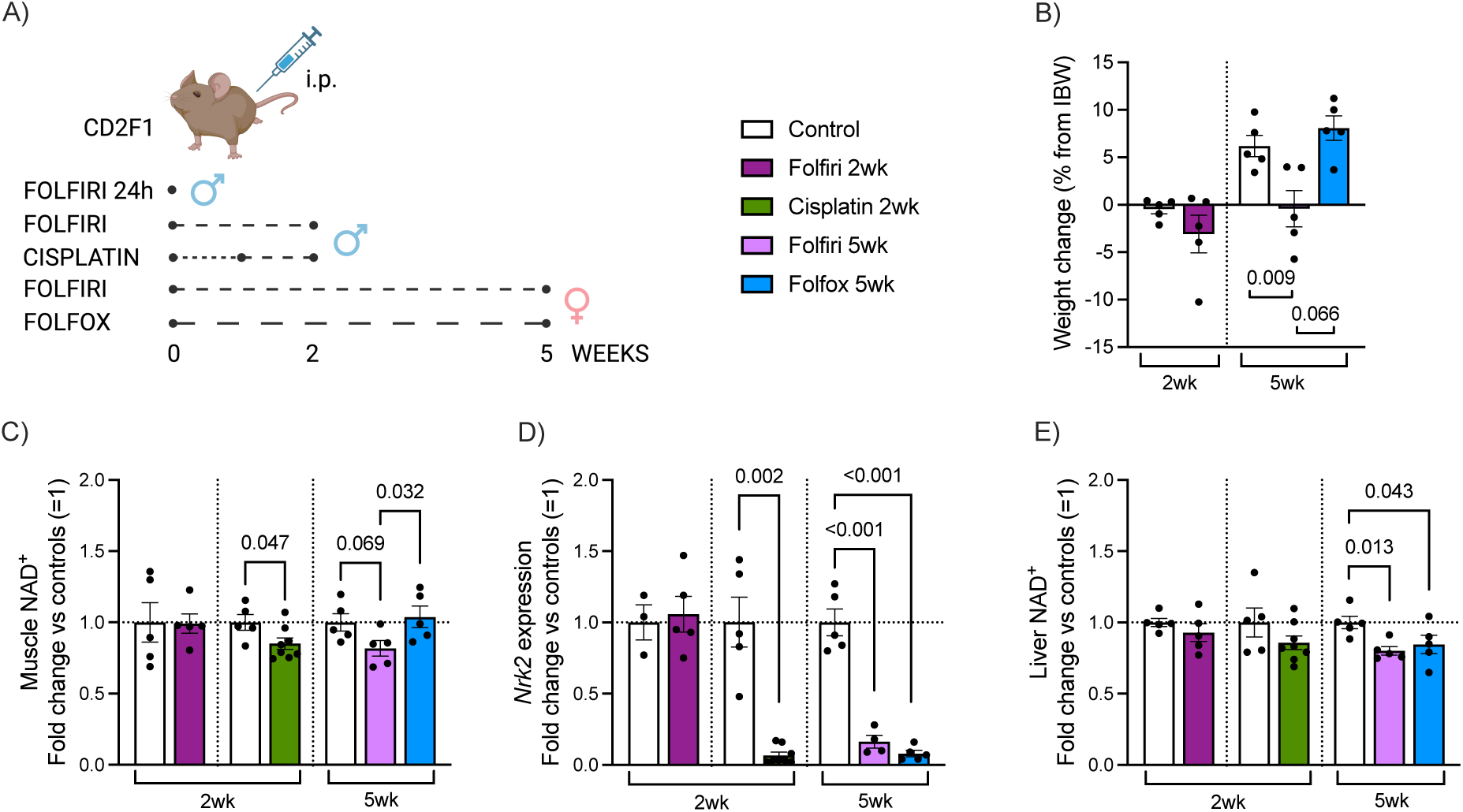
NAD^+^ metabolism alterations in skeletal muscle and liver tissues of chemotherapy-treated mice. A) The study protocol of chemotherapy-treated mice. Male mice were used in the 24-hour (24h) folfiri and the 2-week experiments with cisplatin and folfiri, while females were utilized in the 5-week experiments with folfiri and folfox. The dashed line represents the frequency of chemotherapy administration. B) Weight change (% from initial body weight (IBW)) in the 2-week folfiri (N=5) and the 5-week folfiri (N=5) and folfox (N=5) treated mice and respective controls (N=5) for each group. The weight data from the 2-week cisplatin-treated mice are published by Essex et al (14). C) Quadriceps muscle NAD^+^ content and D) muscle Nrk2 expression in 2-week folfiri (N=5), 2-week cisplatin (N=8) and 5-week folfiri (N=4-5) and folfox (N=5) treated mice and respective controls (N=3-5) for each group. E) Liver NAD^+^ content in 2-week folfiri (N=5), 2-week cisplatin (N=8) and 5-week folfiri (N=5) and folfox (N=5) treated mice and respective controls (N=5) for each group. Data are shown as means ± SEM with individual values. NAD^+^ and Nrk2 data are displayed as fold change versus controls (control mean = 1). Statistical analysis included normality testing with Shapiro Wilk, followed by Student’s t test. Data related to other NAD metabolites, mRNA expression of NAD^+^ biosynthetic and consuming enzymes, and PARP activity are presented in the Supplementary Figures 2-5.

For the cancer time-course experiment, 8-week-old male CD2F1 mice (Inotiv, West Lafayette, IN, USA) were group-housed and randomly assigned to be sacrificed at defined time points following vehicle injection or tumor inoculation (N=5): baseline (day 0) and days 2, 6, 10 and 14. Mice assigned to the tumor group were subcutaneously injected with 1.0 x 10^6^ C26 tumor cells in sterile saline (C26, n = 5) or an isovolumetric subcutaneous injection of vehicle (control, n = 5).

Across all experiments, body weights were recorded daily. At the time of euthanasia, skeletal muscles (gastrocnemius, quadriceps, tibialis anterior) and liver tissue were collected, weighed, snap frozen in liquid nitrogen and stored at –80°C for further biochemical analyses. Chemotherapy regimens and experimental time points were selected based on prior published studies (5,14,15).

### NAD metabolite quantification

NAD metabolites were measured from quadriceps muscle and liver with a slightly modified conventional colorimetric method (16). Metabolite levels were normalized to the total protein content of the sample measured with the Bradford method.

### Real-time quantitative polymerase chain reaction (qRT-PCR)

RNA was extracted from quadriceps muscle and liver with a standard phenol-chloroform method or RNeasy Mini kit (Qiagen, Valencia, CA, USA) following the manufacturer’s protocol. Total RNA was reverse transcribed with Quantitect Reverse Transcription Kit (Qiagen, Hilden, Germany) or Verso cDNA kit (Thermo Fisher Scientific, Waltham, MA, USA). mRNA transcript levels were measured with real-time quantitative PCR using SYBR Green or TaqMan assays (Mm01172899_g1) and normalized to housekeeping genes (Hprt and 36b4, or Tbpl1) using the 2^-ΔΔCT method. SYBR Green primer sequences were as previously described (8).

### PARP activity

PARP activity was measured from quadriceps muscle and liver with the HT Colorimetric PARP/Apoptosis Assay Kit (R&D Systems, Minneapolis, MN, USA) using 2 ug protein per well as per manufacturer’s instructions. Results were normalized to protein concentration of the samples quantified with the Pierce BCA protein assay kit (Thermo Fisher Scientific, Massachusetts, MA, USA).

### Cross-species *skeletal muscle Nrk2/NRK2* expression analyses

Skeletal muscle Nrk2/NRK2 expression in public transcriptomic datasets was analyzed using a combination of web-based differential expression tools, published meta-analysis resources, and custom R workflows, depending on dataset type and data availability (Supplementary Table 1). GEO datasets were queried with GEO2R or, for raw microarray data, normalized and analyzed locally with limma; RNA-seq count matrices were reanalyzed with DESeq2 after filtering and normalization. Summary statistics for datasets without accessible raw expression data were extracted from the original publications or meta-analysis resources (MetaMEx v3.2208; 17). Differentially expressed genes were defined using dataset-specific adjusted p-value and fold-change thresholds as indicated in Supplementary Table 1.

### Statistical analyses

Statistical analyses for experimental data were performed in GraphPad Prism. After normality distribution testing with Shapiro-Wilk, Student’s t test or analysis of variance (ANOVA) was used for normal distribution and Mann-Whitney or Kruskal-Wallis for non-normal distribution. These were followed by post hoc analyses for planned comparisons: Fisher’s Least Significant Difference (LSD) test for ANOVA and Uncorrected Dunn’s test for Kruskal-Wallis. The associations between variables in the correlation analyses were tested with either Pearson correlation for normally distributed data or Spearman’s rank correlation coefficient for nonparametric data. The statistical significance threshold was set at *p*<0.05.

## Results

### Skeletal muscle NAD^+^ metabolism is disturbed by chemotherapeutic agents upon muscle atrophy development

We first characterized how chemotherapy alters skeletal muscle NAD^+^ metabolism in healthy mice using regimens known to promote muscle wasting (5,15). We analyzed tissue samples from *in vivo* experiments involving cisplatin, folfiri and folfox, including previously reported cisplatin cohort (14). The study designs are summarized in Figure 1A.

In an initial experiment, healthy wild-type male mice were treated with either folfiri or cisplatin for up to two weeks. Folfiri did not affect body weight or muscle mass (Figs. 1B, S1A,B), whereas cisplatin induced a decrease in body weight and muscle loss (14). Because short-term (2 weeks) folfiri exposure did not trigger weight loss, we extended our analyses to a 5-week timepoint and included an additional folfox-treated group, previously used to explore NAD^+^ metabolism in cancer- and chemotherapy-induced cachexia (8). Analyses of available samples from female mice revealed reduced final body weight and muscle mass in folfiri-treated animals compared to controls (Figs. 1B, S1C,D). In contrast, folfox-treated mice maintained stable body weight but exhibited mild muscle loss (Figs. 1B, S1C,D). Overall, cisplatin induced the most severe cachexia phenotype, followed by prolonged folfiri exposure, which resulted only minor effects.

Muscle NAD metabolite measurements revealed distinct, treatment-dependent effects. Consistent with preserved muscle mass and body weight, 2-week folfiri did not alter NAD metabolites (Figs. 1C, S2A) or their ratios (Fig. S2B). In contrast, cisplatin reduced muscle NAD^+^ (−14%) (Fig. 1C) along with declines in NADH and NADPH (Fig. S2C), in line with the severe body and muscle weight loss. While the NAD^+^/NADH ratio remained unchanged, indicating proportional depletion of both species, the NADP^+^/NADPH ratio increased (Fig S2D), suggesting a shift toward a more oxidized cellular state. The 5-week folfiri elicited a near-significant decrease (−18%) in muscle NAD^+^ levels (Fig. 1C) and a reduction in NADP^+^ (Fig. S2E), aligning with the modest declines in body and muscle mass. Both NAD^+^/NADH and NADP^+^/NADPH were significantly decreased (Fig. S2F) indicating a shift toward a more reduced intracellular redox environment. In contrast, the 5-week folfox treatment did not alter NAD metabolites or their ratios (Figs. 1C, S2E,F), in line with the absence of overt muscle atrophy. Altogether, these findings indicate that the loss of skeletal muscle NAD^+^ associates with the development and the severity of chemotherapy-induced muscle atrophy.

To investigate the mechanisms underlying skeletal muscle NAD^+^ decline, we assessed the mRNA expression of key enzymes involved in NAD^+^ biosynthetic pathways. In skeletal muscle, NAD^+^ is primarily generated via the salvage pathway through *Nampt* (18). Consistent with preserved muscle NAD^+^ levels, the 2-week folfiri treatment did not alter the expression of muscle NAD^+^ biosynthetic enzymes (Fig. S3A). In contrast, cisplatin-treated mice showed no change in *Nampt*, but exhibited reduced levels of the other salvage pathway enzymes *Nmnat 1* and *Nmnat 3*, along with a marked downregulation of *Nrk2* (Figs. 1D, S3B). We also observed suppression of enzymes within the Preiss-Handler (PH) pathway, including *Naprt* (Fig. S3B), although this pathway is considered minor in basal NAD^+^ maintenance in skeletal muscle (19).

At the 5-week timepoint, folfiri-treated mice selectively downregulated muscle *Nrk2*, with no significant changes in other NAD^+^ biosynthetic enzymes (Figs. 1D, S3C), consistent with the modest decline in muscle NAD^+^. Notably, folfox reduced muscle *Nrk2* expression despite preserved NAD^+^ levels, which was accompanied by upregulation of the ubiquitously expressed isoform *Nrk1* (20) (Figs. 1D, S3C). Consistently, muscle *Nrk2* downregulation was detectable as early as 24 hours after folfiri treatment, preceding any measurable muscle loss (Fig. S3D). Collectively, these findings identify *Nrk2* as an early, chemotherapy-responsive NAD^+^ biosynthetic gene, with its suppression preceding NAD^+^ loss and muscle wasting.

Finally, we assessed the skeletal muscle mRNA expression of NAD^+^-consuming enzymes and the activity of the main NAD^+^ consumers, PARPs. Chemotherapy had minimal effects on the expression of NAD^+^-consuming enzymes and did not alter PARP activity at either timepoint (Fig. S3A,B,E). Overall, these findings suggest that chemotherapy does not alter skeletal muscle NAD^+^ consumption in healthy mice.

### Prolonged chemotherapy alters hepatic NAD^+^ metabolism

Given that chemotherapy affects multiple tissues, we next investigated hepatic NAD^+^ metabolism, which is disrupted in cancer-induced cachexia (8). Neither folfiri nor cisplatin impacted hepatic NAD^+^ levels when administered for 2 weeks (Figs. 1E, S4A,C). However, folfiri reduced NADP^+^ levels without affecting metabolite ratios (Fig. S4A,B). In contrast, cisplatin-treated mice exhibited a non-significant increase in NADH that nonetheless resulted in a lower NAD^+^/NADH ratio (Fig. S4C,D), suggestive of a shift in hepatic redox balance. At the 5-week timepoint, both folfiri and folfox regimens induced modest but significant decreases in hepatic NAD^+^ levels (−15 to −20%) with no corresponding changes in other metabolites or their ratios (Figs. 1E, S4E,F). Overall, these findings indicate that hepatic NAD^+^ loss occurs primarily with prolonged chemotherapy exposure, implying a limited metabolic impact in healthy mice.

To address underlying mechanisms, we analyzed the mRNA expression of enzymes in the NAD^+^ biosynthetic pathways. In the liver, NAD^+^ is predominantly generated via the *de novo* pathway from tryptophan (21). Consistent with stable NAD^+^ levels after 2 weeks of folfiri, only modest transcriptional changes were observed, including trends toward increased *Qprt* and reduced salvage pathway enzyme *Nrk1* (Fig. S5A). In cisplatin-treated animals, the hepatic *de novo* pathway genes were reduced, with a trend toward suppression of the PH pathway, while the salvage pathway enzymes *Nampt* and *Nrk1* were upregulated (Fig. S5B). In contrast, under conditions of modest NAD^+^ loss at 5 weeks, folfiri mainly reduced hepatic *Nrk1*, whereas folfox induced broader alterations in NAD^+^ biosynthesis, including increased *Tdo* and *Nampt* and decreased *Naprt* (Fig. S5C). These findings highlight treatment-specific reorganization of hepatic NAD^+^ biosynthesis, which may contribute to the observed decrease in NAD^+^ levels.

Finally, we assessed whether hepatic NAD^+^ consumption contributes to the observed metabolic changes. The expression of NAD^+^-consuming enzymes was largely unchanged across conditions (Fig. S5A,B,C). Similarly, hepatic PARP activity remained stable, with only minor decline observed in cisplatin-treated mice (Fig. S5D). Collectively, these findings suggest that chemotherapy-induced hepatic NAD^+^ loss is unlikely to be driven by increased NAD^+^ consumption, and instead may primarily reflect altered NAD^+^ biosynthetic capacity.

### Skeletal muscle *Nrk2* expression is altered in conditions influencing muscle mass in rodents and humans

Given the growing evidence that skeletal muscle NAD^+^ depletion and downregulation of *Nrk2* are associated with cachexia (8), we asked whether these changes reflect ongoing muscle atrophy. We therefore examined the relationship between muscle mass at time of euthanasia and muscle NAD^+^ levels or *Nrk2* expression in chemotherapy-treated mice exhibiting ongoing or developing muscle wasting (2-week cisplatin and 5-week folfiri and folfox) and our previously published cancer cachexia datasets, including tumor-bearing mice with chemotherapy-naive or folfox-treated conditions (8).

When individual experiments were analyzed separately, muscle NAD^+^ showed no consistent relationship with muscle mass, with substantial variability across models and signs of dataset specific effects, particularly in more severe conditions (Fig. S6A–E). In contrast, muscle *Nrk2* expression was positively associated with muscle mass in a subset of models (Fig. S6F-J), but not uniformly in all experimental groups, indicating context-dependent variability.

When all datasets were visualized together, muscle NAD^+^ displayed an overall but variable relationship with muscle mass, with clear differences between models (Fig. 2A,C). In comparison, muscle *Nrk2* expression showed more apparent positive association across models, although the strength of this relationship remained model-dependent (Fig. 2B,D). For both muscle NAD^+^ and *Nrk2*, these associations appeared more evident in tumor-bearing models than in chemotherapy-treated healthy mice, consistent with more pronounced metabolic adaptations in the presence of a tumor.

**Figure 2.**
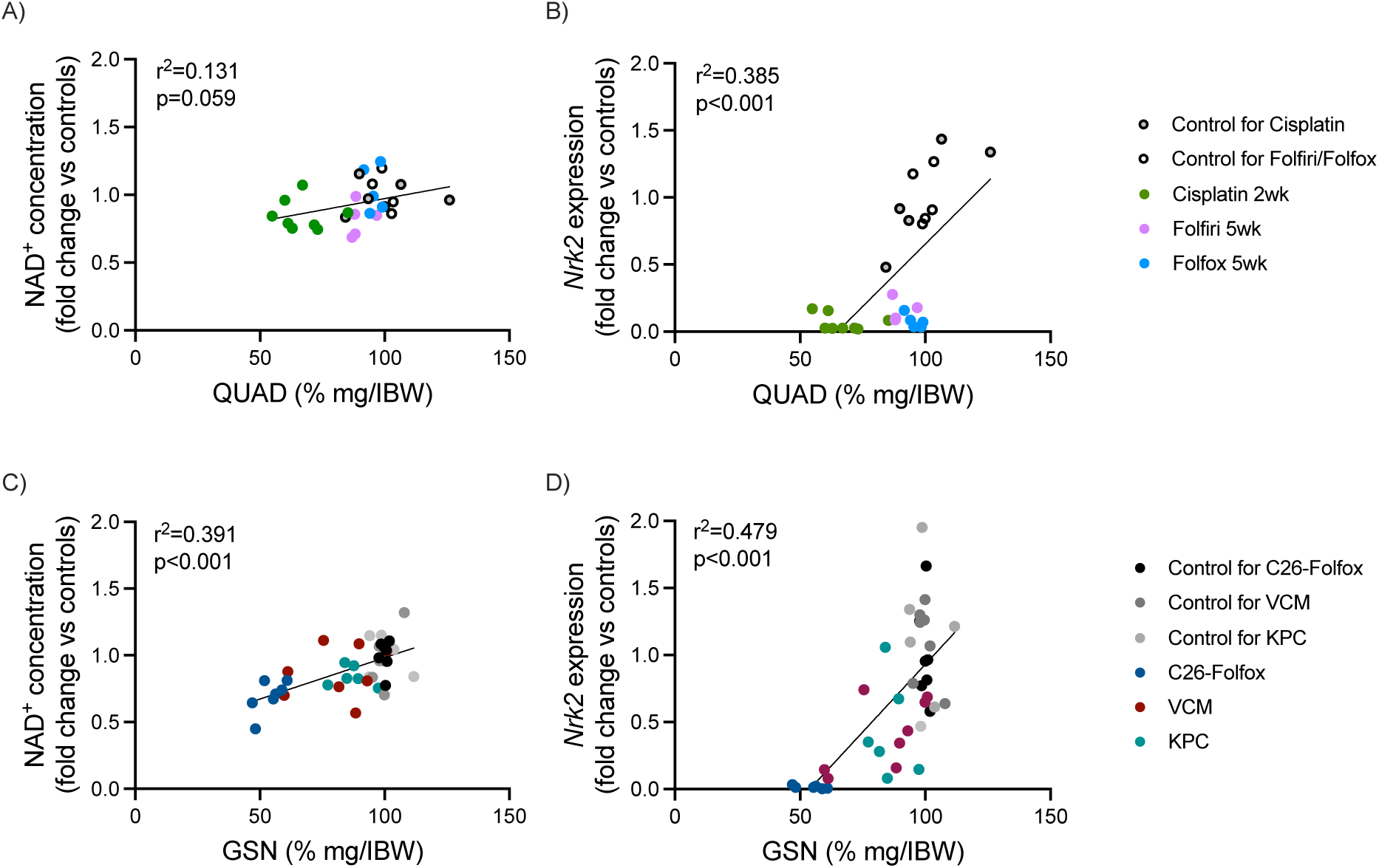
The relationship of skeletal muscle NAD^+^ levels and Nrk2 expression with muscle mass across chemotherapy and cancer cachexia mouse models. Scatter plots showing the association between quadriceps (QUAD) muscle mass and A) NAD^+^ levels and B) Nrk2 expression in the combined dataset of 2-week cisplatin-treated and 5-week folfiri- and folfox-treated mice and their controls. Corresponding associations are shown for gastrocnemius (GSN) muscle mass and C) NAD^+^ levels, and D) Nrk2 expression in the combined dataset of C26-Folfox, Villin-Cre/Msh2 (VCM) and KPC mice and their respective controls (8). The combined muscle NAD^+^ and Nrk2 data are displayed as fold change versus controls (control mean = 1). Muscle mass is reported as wet weight normalized to IBW and represented as percentage of the mean of the respective control group. The distribution of data was tested with Shapiro Wilk and the associations between variables were tested with either Pearson correlation for normally distributed data or Spearman’s rank correlation coefficient for nonparametric data. Correlations for individual experiments are presented in Supplementary Figure 6.

Collectively, these observations suggest that skeletal muscle NAD^+^ levels and *Nrk2* expression track changes in muscle mass in a context-dependent manner. While neither readout exhibited a uniform relationship across all models, the more frequent associations observed for *Nrk2* expression across datasets supports its potential utility as an indicator of muscle atrophy in wasting conditions.

These findings prompted us to seek additional evidence for skeletal muscle *Nrk2* expression in muscle wasting. Given the limited transcriptomic data available for chemotherapy-induced cachexia, we turned to the broader cancer cachexia literature. A recent meta-analysis by Zhao et al (17), encompassing 13 studies across multiple mouse models, including Colon-26 carcinoma (C26), KPC pancreatic cancer, Lewis lung carcinoma (LLC), and genetically engineered mouse models of pancreatic cancer (KPP and KPC GEMM), identified *Nrk2* as consistently downregulated in skeletal muscle across all cancer cachexia models, with the most pronounced suppression observed in the C26 model (Fig. 3A). Although temporal data remains limited, analysis of our previously published C26 model dataset showed that muscle *Nrk2* expression declined during cachexia development, becoming reduced from day 6 onward (Fig. S6K), with transient fluctuations reflecting variability in disease progression and metabolic alterations (22). Conversely, in our previous study, muscle *Nrk2* expression was shown to be restored by activin receptor ligand blockade (7), which is consistent with data from multiple cachexia-targeted therapies that increase muscle mass (Fig. 3B) (23–25 (GSE56555; GSE214603; GSE123310)). Together, these findings support skeletal muscle *Nrk2* downregulation as an early, dynamic, and therapeutically reversible feature of cancer cachexia in rodents, which may be modulated by muscle mass restoring interventions.

**Figure 3.**
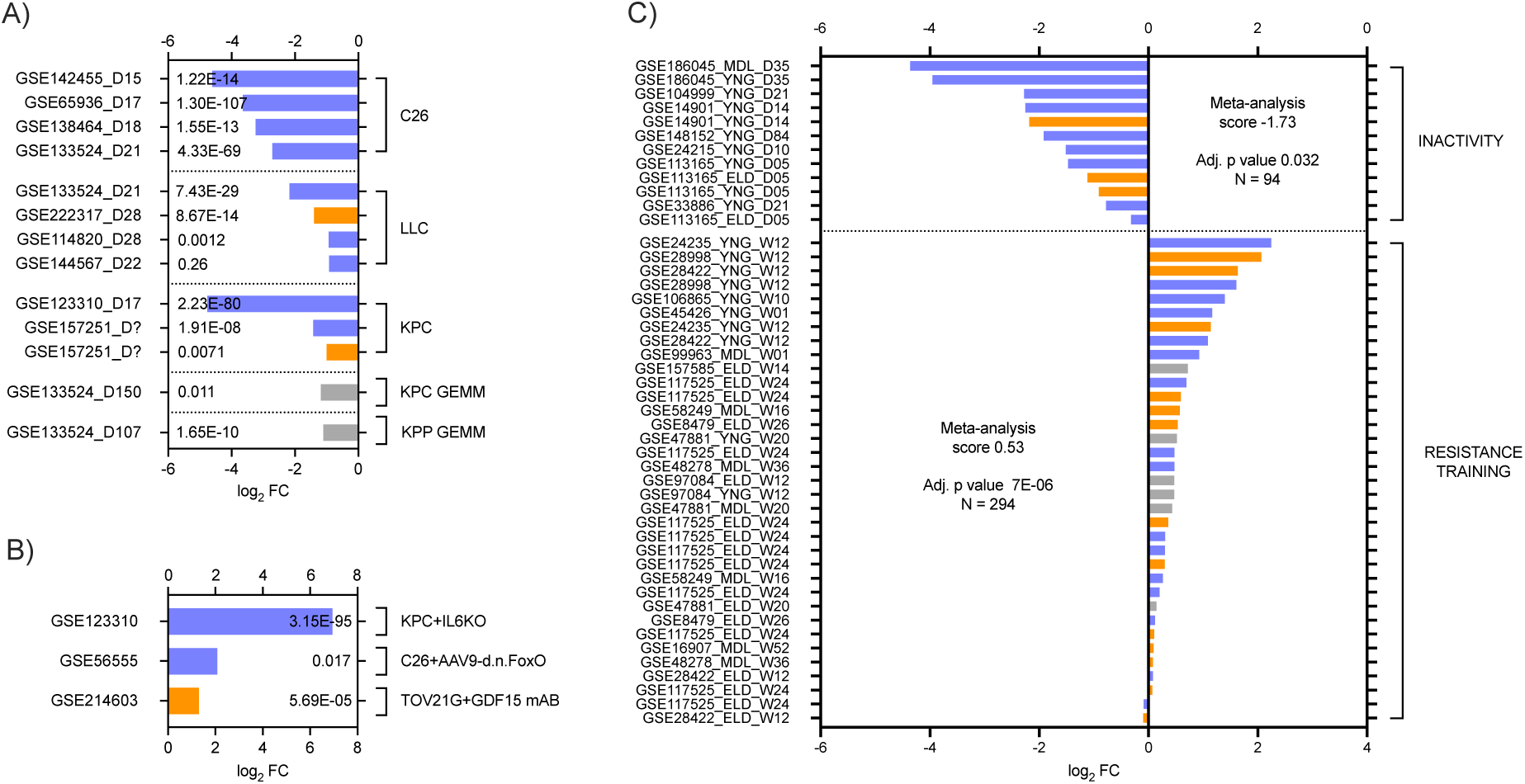
Skeletal muscle Nrk2 expression across rodent cancer cachexia models, cachexia-targeted therapies, and human muscle plasticity conditions. A) Skeletal muscle Nrk2 expression in rodent cancer cachexia models including C26 (N=3-6/group), LLC (N=4-8/group), and pancreatic cancer cachexia models (KPC, KPC GEMM, KPP GEMM) (N=3-4/group) expressed as log_2_FC compared to each experiment’s control group. FDR adjusted p values for each study are reported inside the frame. Data were collected from the meta-analysis of Zhao et al (17): Nrk2 DEG fold change and FDR values were retrieved from Table S2 and FC transformed to log_2_FC. The duration of the experiment is stated as days (D) after the GEO identification number. Two studies did not report the specific duration of the experiment (marked as “D?”). B) Skeletal muscle Nrk2 expression following cachexia-targeted interventions in three mouse models: KPC pancreatic cancer with IL-6-deficient tumors (KPC-IL6KO vs KPC, n=4/group), C26 colon cancer with AAV9-mediated inhibition of FoxO-dependent transcription (C26-d.n.FoxO vs C26-empty vector, n=4 vs n=4/group), and TOV21G ovarian cancer with anti-GDF15 antibody treatment (mAB2 vs IgG, n=11 vs n=14). Bars show log2 fold change of the intervention group versus the untreated cachexia group; FDR-adjusted p-values are shown inside each bar. C) Skeletal muscle NRK2 expression in humans following inactivity (N=2-12/ group) or undergoing resistance training (N=2-24/experimental group) expressed as log_2_FC and p values adjusted for multiple comparison with the Bonferroni method. Inactivity studies include every study lasting five days or more. Each study is listed with its GEO identification number, age of the studied cohort (YNG = young, MDL = middle aged, ELD = elderly) and duration of the experiment in days (D) or weeks (W). Data were collected from the MetaMEx database (accessed 5^th^ of March 2025). Blue = male, orange = female, grey = unknown or both sexes.

We next assessed skeletal muscle *NRK2* expression in human muscle-wasting conditions. In a study of cachectic pancreatic cancer patients (26), muscle *NRK2* ranked among the top 20 most differentially expressed transcripts as compared to healthy controls (Table 1). In additional datasets, we observed modest but non-significant reductions (27,28) or no differences in muscle *NRK2* expression between cancer patients and controls (29) (Table 1). However, these datasets are limited by small cohorts, lack of non-cancer controls, and incomplete cachexia stratification. We further found reduced *NRK2* expression in skeletal muscle of elderly sarcopenic patients (30) and patients with critical illness myopathy (31) compared to healthy counterparts (Table 1). Collectively, these observations suggest that skeletal muscle *Nrk2* expression decreases across diverse atrophic conditions in humans.

**Table 1.** Skeletal muscle *NRK2* expression across human muscle-wasting cohorts.

| Disease | Publication | GEO accession | Patients | Log<sub>2</sub>FC (patient vs control) | Adjusted p value | Additional information |
| --- | --- | --- | --- | --- | --- | --- |
| Cancer | Talbert et al (2019) | GSE133523 | controls with nonmalignant disease (N=5)<br>PDAC (N=5) | -1.912 | 0.0002 | cancer group weight change mean 10.8% ± 1.0 |
|  | Narasimhan et al (2021) | GSE133979 | controls with nonmalignant disease (N=11)<br>PDAC (N=23) | -0.367 | 0.800 | cancer group weight loss grade 0-4 |
|  | Gallagher et al (2012) | GSE34111 | controls with nonmalignant disease (N=5)<br>upper gastrointestinal cancer (N=12) | -0.561 | 0.342 | cancer group weight change -25% - +4% |
|  | Stephens et al (2010) | GSE18832 | controls with nonmalignant disease (N=3)<br>upper gastrointestinal cancer (N=18)" | 0.052 | 0.999 | cancer group weight change -22% - +0.5% |
| Sarcopenia | Zuo et al (2025) | GSE226151 | healthy aged (HA) (N=20)<br>presarcopenia (PS) (N=19)<br>sarcopenia (S) (N=19) | S vs HA -1.277<br>S vs PS -1.014<br>PS vs HA -0.264 | S vs HA 0.035<br>S vs PS 0.199<br>PS vs HA 0.759 | presarcopenia = low muscle strength but unchanged mass |
| Critical illness myopathy | Llano-Diez et al (2019) | - (data from publication) | healthy controls (N=6)<br>critical illness myopathy patients (N=7) | -2.047 | 0.0142 |  |

*Per-cohort log<sub>2</sub> fold change and Benjamini–Hochberg adjusted p-value for the patient-versus-control contrast; see Methods for reanalysis pipeline. Where indicated, values were taken from the source publication. The Additional information column summarizes clinical features of weight loss or muscle atrophy as report. PDAC = pancreatic ductal adenocarcinoma.*

To investigate skeletal muscle *NRK2* regulation under conditions that affect muscle mass, we analyzed human inactivity and resistance training studies. Intriguingly, skeletal muscle *NRK2* expression was strongly decreased upon inactivity caused by bed rest or limb immobilization (Fig. 3C). Conversely, resistance training, a robust inducer of muscle hypertrophy (32) significantly upregulated muscle *NRK2* expression (Fig. 3C). In summary, these findings show that *NRK2* expression in human skeletal muscle decreases with disuse and increases with hypertrophy.

## Discussion

NAD^+^ deficiency has been previously implicated in the pathophysiology of experimental cancer-induced cachexia (7,8). Here, we extend this concept by demonstrating that chemotherapy alone can act as a regulator of NAD^+^ metabolism, independent of tumor burden. Specifically, we show that chemotherapeutic agents lower skeletal muscle NAD^+^ levels and reduce *Nrk2* expression, two key features previously associated with cancer cachexia (7,8). By further identifying consistent muscle *Nrk2* transcriptional responses across multiple muscle wasting and hypertrophy conditions in rodent models and humans, we highlight muscle *Nrk2* expression as a conserved feature associated with muscle mass regulation across species, prompting to design targeted interventions for muscle atrophy. Together, our findings underscore the potential of NAD^+^ metabolism as a diagnostic and therapeutic target for cancer- and chemotherapy-induced cachexia.

This work represents the first investigation of skeletal muscle NAD^+^ metabolism following chemotherapy treatment, particularly in the context of chemotherapy-induced muscle wasting. Consistent with our previous findings (7,8), we observed that muscle NAD^+^ loss parallels muscle atrophy in chemotherapy treated wild-type mice. Cisplatin induced the most pronounced reductions in both NAD^+^ levels and muscle mass, whereas prolonged folfiri treatment resulted in more modest declines. Although the extent of NAD^+^ loss was less pronounced (−14 to −18%) than in tumor-bearing cachectic mice (−13 to −31%) (8), this pattern reinforces the notion that chemotherapeutic agents alone can have negative impact on muscle NAD^+^ homeostasis. These findings align with prior evidence of chemotherapy-induced muscle mitochondrial dysfunction and stress responses (5), processes that are closely linked to NAD^+^ metabolism, further supporting a role for NAD^+^ metabolism in muscle wasting and cachexia.

In contrast to skeletal muscle, the liver exhibited a more modest response to chemotherapy in healthy mice. While prior work has shown marked hepatic NAD^+^ depletion in tumor-bearing models (8), prolonged chemotherapy exposure in healthy mice induced only mild reductions in hepatic NAD^+^ levels. These hepatic changes were substantially smaller (−15 to −20%) as compared to those observed in cachectic tumor hosts receiving chemotherapy (−31 to −60%) (8), which might reflect lower degree of systemic stress and injury or stronger adaptation capacity and metabolic flexibility in healthy mice. Together with the skeletal muscle findings, the data implies that tumor burden amplifies chemotherapy-induced metabolic disturbances, supporting a cumulative or interacting effect on tissue NAD^+^ homeostasis. This divergence between chemotherapy-treated healthy mice and tumor-bearing conditions highlights tissue-specific vulnerability to chemotherapy-induced metabolic defects and differential sensitivity to chemotherapeutic stress in diseased and healthy mice. However, the mechanisms by which liver and skeletal muscle coordinate systemic NAD^+^ metabolism remain incompletely understood and warrant further investigation.

Mechanistically, our data indicate that chemotherapy-induced muscle NAD^+^ loss is primarily driven by impaired NAD^+^ biosynthesis, rather than increased consumption. A consistent transcriptional response across chemotherapy treatments was the downregulation of muscle *Nrk2*. Prior studies suggest that Nrk2 can be rate-limiting for NAD^+^ biosynthesis during regeneration (33) or supportive during stress (9,34). Notably, muscle *Nrk2* downregulation occurred upon prolonged folfox treatment, despite preserved NAD^+^ levels, suggesting that *Nrk2* repression may precede NAD^+^ depletion or become functionally uncoupled from NAD^+^ biosynthesis. Potential upstream regulators of *Nrk2* include PPARα, a known transactivator (9), cytokine-driven inflammation (35), or chemotherapy-induced activin signaling (36), the latter of which is reversed by activin inhibition (7). In the liver, mild NAD^+^ loss upon prolonged folfiri and folfox treatment was accompanied by modest downregulation of some PH and salvage pathway gene expression and increased *de novo* pathway gene *Tdo,* suggesting a reorganization of NAD^+^ metabolism in which increased *de novo* pathway activity is insufficient to fully maintain NAD^+^ levels. Nonetheless, the precise drivers of tissue-specific NAD^+^ dysregulation during chemotherapy treatment remain to be defined.

Previous work has shown that mRNA levels of muscle NAD^+^ biosynthetic enzymes *Nampt* and *Nmnat3* positively correlate with muscle mass in mice (37), supporting a link between NAD^+^ metabolism and muscle preservation or atrophy. Building on this, we found that muscle *Nrk2* expression appeared to show a more consistent association with muscle mass than NAD^+^ levels, across healthy mice and in cancer- and chemotherapy-induced muscle wasting. Converging evidence from multiple mouse models further indicates reduced muscle *Nrk2* expression as an early feature of muscle atrophy as demonstrated by acute doxorubicin exposure (13), progressive decline during C26-induced cachexia and early suppression in the diaphragm of KPC mice prior to overt muscle loss (38). This pattern is further supported by integrated analyses showing consistent *Nrk2* suppression across cancer cachexia mouse models (39). Together, these findings support muscle *Nrk2* downregulation as an early and consistent feature of muscle atrophy across experimental models.

Extending these observations to humans, public transcriptomic datasets (30,31, MetaMEx) suggest that skeletal muscle *NRK2* expression is reduced across atrophic conditions and inactivity while hypertrophic stimuli such as resistance training increase its expression in humans. Accordingly, muscle *NRK2* was identified among the top-most downregulated genes after disuse, and among the most upregulated gene after recovery from atrophy in humans (40). In line with this bidirectional regulation, several independent cachexia-targeted interventions that preserve or increase in muscle mass, including activin receptor ligand blockade, IL-6 deficiency, inhibition of FoxO signaling, or neutralization of GDF15 (7,23–25), are accompanied by increased muscle *Nrk2* expression. Although *Nrk1/2* double knockout mice do not develop muscle atrophy (33), arguing against a causal role in muscle catabolism, these observations collectively support a model in which muscle *Nrk2* expression dynamically tracks changes in muscle mass, reflecting transitions between catabolic and anabolic muscle states rather than a disease-specific response. Altogether, our findings suggest that muscle *Nrk2* expression could serve as a predictor of muscle mass or atrophy, with potential utility for monitoring treatment responses over time.

Several study limitations should be acknowledged. The relatively small sample size in the mouse experiments may limit the statistical power, particularly for detecting subtle changes in NAD^+^ metabolites and muscle *Nrk2* expression. Additionally, the lack of detailed temporal analysis prevents us from defining the dynamic progression of NAD^+^ during the development of muscle atrophy. Sex-specific differences also remain incompletely addressed, as short- and long-term experiments were conducted in male and female mice, respectively. Despite these limitations, previous work demonstrating the efficacy of NAD^+^ repletion strategies across sexes (8) supports the broader relevance of our findings. Future studies should incorporate longitudinal designs, balanced sex representation, and mechanistic investigations to define upstream regulators of *Nrk2* and NAD^+^ metabolism.

In conclusion, we demonstrate that chemotherapy-induced cachexia shares a core NAD^+^ metabolic signature with cancer cachexia, characterized by skeletal muscle NAD^+^ loss and *Nrk2* downregulation. These findings underscore the therapeutic potential of targeting NAD^+^ metabolism to combat muscle loss during chemotherapy. Furthermore, we identify muscle *Nrk2* expression as a conserved and responsive marker of muscle mass changes, with potential utility for monitoring disease progression and treatment efficacy. Understanding the mechanisms governing tissue-specific NAD^+^ regulation and *Nrk2* expression dynamics will be essential for translating these findings into clinical strategies aimed at mitigating cachexia and preserving metabolic health in cancer patients.

## Conflict of interest

The authors declare no conflict of interest.

## Funding

This study was supported by research grants from The Finnish Medical Foundation, The Paulo Foundation, The Maud Kuistila Memorial Foundation, The Orion Research Foundation and The Ida Montin Foundation to N.P., by Fondazione AIRC (IG 2024—ID. 31126 project, PI: Fabio Penna) to F.P., the Department of Pathology and Cancer Center at University of Colorado Anschutz, NIH/NIAMS (R01AR079379, R01AR080051) and Wings of Hope for Pancreatic Cancer Research to A.B., and the Cancer Foundation Finland and the Research Council of Finland Profi6 funding (336449) awarded to the University of Oulu to E.P..

## Code availability

Indicated analyses were performed in R (v4.5.3); code is private during review at https://github.com/gammon-bio/nrk2_analysis and will be made public with a Zenodo DOI on acceptance.

## Notes

### Competing Interest Statement

The authors have declared no competing interest.

