## Supplementary materials for "Chemotherapy induces tissue NAD^+^ loss, and downregulation of NAD^+^ biosynthetic enzyme *Nrk2* marks muscle wasting"

***Supplementary Table 1.***

| Dataset / Accession | Source | Disease / model (intervention) | Species | Tissue | Platform / data type | Analysis method | Contrast(s) | Correction | Notes |
|---|---|---|---|---|---|---|---|---|---|
| Zhao et al. (2024) | Published meta-analysis (their Table S2) | Multiple rodent cachexia models | Rodent | Multiple | RNA-seq (RPKM) | Extracted from publication; FC converted to log2FC | As reported per study | FDR (as reported) | Per-study values drawn from Zhao et al. Table S2 |
| GSE214603 | GEO | TOV21g ovarian cancer (GDF15-neutralizing antibody) | Mouse | GSN | RNA-seq (raw counts) | DESeq2 [PMID: 25516281] (median-of-ratios, Wald, BH) | cachectic vs control; intervention vs control; intervention vs cachectic | Benjamini-Hochberg | log2FC and adjusted p graphed |
| GSE123310 | GEO | KPC pancreatic cancer (+/- tumor-derived IL-6) | Mouse | QUAD | RNA-seq (raw counts) | DESeq2 [PMID: 25516281] (median-of-ratios, Wald, BH) | cachectic vs control; intervention vs control; intervention vs cachectic | Benjamini-Hochberg | log2FC and adjusted p graphed |
| GSE56555 | GEO | C26 colon cancer (AAV9 inhibition of FoxO-dependent transcription) | Mouse | TA | Microarray (Affymetrix Mouse Gene 1.0 ST) | GEO2R [PMID: 23193258] (limma, default settings) | C26-empty vector (n=4) vs C26-d.n.FoxO (n=4) | Benjamini-Hochberg | All C26 samples retained; outlier excluded in original publication could not be unambiguously identified from GEO sample annotation |
| MetaMEx | Meta-analysis web app (accessed 5 Mar 2025) | Human inactivity and resistance training | Human | SM | Aggregated meta-analysis output | Nrk2/NRK2-specific log2FC and adjusted p extracted from MetaMEx output | Per study | Adjusted p (as reported) | Inactivity studies restricted to >=5 days |
| GSE133979 (Narasimhan et al. 2021) | GEO | Pancreatic ductal adenocarcinoma (PDAC) | Human | RA | RNA-seq (raw counts) | DESeq2 [PMID: 25516281] (median-of-ratios, Wald, BH) | PDAC (n=23) vs nonmalignant-disease control (n=11) | Benjamini-Hochberg | cancer group weight-loss grade 0-4 |
| GSE133523 (Talbert et al. 2019) | GEO | Pancreatic ductal adenocarcinoma (PDAC) | Human | RA | RNA-seq (raw counts) | DESeq2 [PMID: 25516281] (median-of-ratios, Wald, BH) | PDAC (n=5) vs nonmalignant-disease control (n=5) | Benjamini-Hochberg | cancer group mean weight change 10.8% +/- 1.0 |
| GSE226151 (Zuo et al. 2025) | GEO | Age-related sarcopenia | Human | SM (not specified) | RNA-seq (raw counts) | DESeq2 [PMID: 25516281] (median-of-ratios, Wald, BH) | sarcopenia (n=19) vs healthy aged (n=20); sarcopenia (n=19) vs presarcopenia (n=19); presarcopenia (n=19) vs healthy aged (n=20) | Benjamini-Hochberg | presarcopenia = low muscle strength with unchanged mass |
| GSE34111 (Gallagher et al. 2012) | GEO | Upper gastrointestinal cancer | Human | QUAD | Microarray (raw CEL; Affymetrix | RMA via affy [PMID: 14960456] + limma [PMID: 25605792] | upper GI cancer (n=12) vs nonmalignant-disease control (n=5) | Benjamini-Hochberg | Annotation via hgu133plus2.db; highest-mean-expression probe |

| | | | | | | | | | |
|---|---|---|---|---|---|---|---|---|---|
| | | | | | HG-U133 Plus 2.0) | (empirical Bayes, BH) | | | retained per gene; cancer group weight change -25% to +4% |
| GSE18832 (Stephens et al. 2010) | GEO | Upper gastrointestinal cancer | Human | RA | Microarray (raw CEL; Affymetrix HG-U133 Plus 2.0) | RMA via affy [PMID: 14960456] + limma [PMID: 25605792] (empirical Bayes, BH) | upper GI cancer (n=18) vs nonmalignant-disease control (n=3) | Benjamini-Hochberg | Annotation via hgu133plus2.db; highest-mean-expression probe retained per gene; cancer group weight change -22% to +0.5% |
| Llano-Diez et al. (2019) | Published supplementary data (no GEO accession) | Critical illness myopathy | Human | TA | Raw data not publicly available | Transcribed from publication | critical illness myopathy patients (n=7) vs healthy controls (n=6) | As reported | log2FC and adjusted p transcribed from supplementary data |

*External transcriptomic datasets and Nrk2/NRK2 analysis details. Public datasets used to assess Nrk2/NRK2 expression across cachexia, sarcopenia, and muscle inactivity, with the source, disease or model, species, tissue, platform, analysis method, and contrast(s) for each. Raw RNA-seq count matrices were reanalyzed with DESeq2 and raw microarray data with RMA/limma or GEO2R; where raw data were unavailable, summary statistics were taken from the original publications or meta-analysis resources. PMIDs for analysis tools are given in the Analysis method column. GSN, gastrocnemius; QUAD, quadriceps; TA, tibialis anterior; SM, skeletal muscle; RA, rectus abdominis; BH, Benjamini-Hochberg; FDR, false discovery rate.*

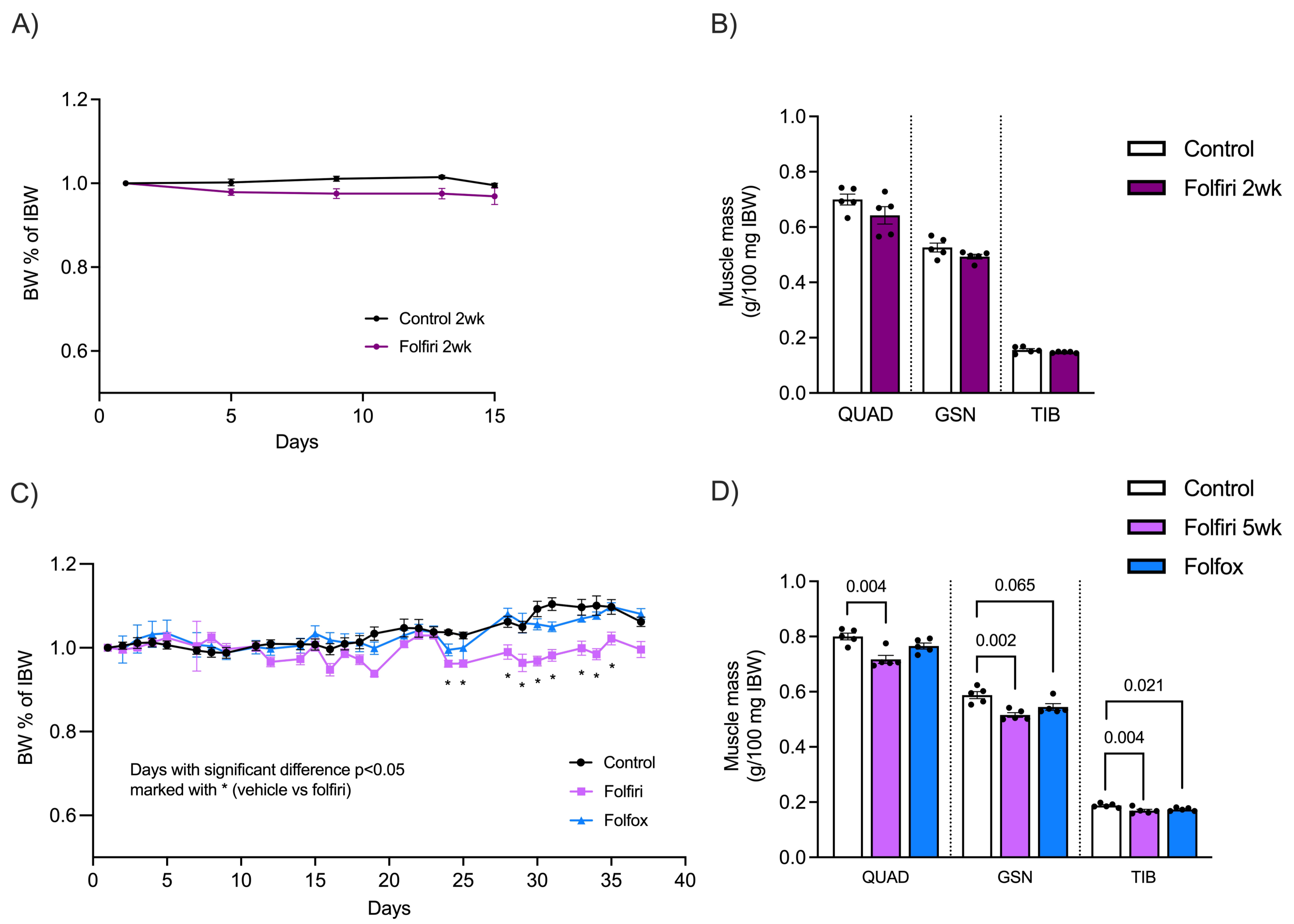


***Supplementary figure 1**. **Body weight development and muscle masses in the 2- and 5-week chemotherapy experiments, related to Figure 1**. Data from the 2-week cisplatin experiment have been published before (14).*

*A) Body weight (BW) development (% from initial body weight (IBW)) in the 2-week folfiri experiment (N=5 per group).*

*B) Quadriceps (QUAD), gastrocnemius (GSN) and tibialis (TIB) muscle masses in the 2-week folfiri experiment (N=5 per group).*

*C) Body weight (BW) development (% from initial body weight (IBW)) in the 5-week folfiri and folfox experiment (N=5 per group).*

*D) Quadriceps (QUAD), gastrocnemius (GSN) and tibialis (TIB) muscle masses in 5-week folfiri and folfox experiment (N=5 per group).*

*Weight development data are shown as mean ± SEM, whereas muscle mass is presented as individual values with mean ± SEM. Statistical analysis for weight development was performed with two way repeated measures ANOVA with time and group for factors followed by Tukey's post hoc multiple comparisons test. Statistical analysis for muscle masses was performed using two-tailed Student's t test or one-way ANOVA followed by Fisher's LSD test for parametric data, and Mann-Whitney or Kruskal-Wallis followed by uncorrected Dunn's test for nonparametric data.*

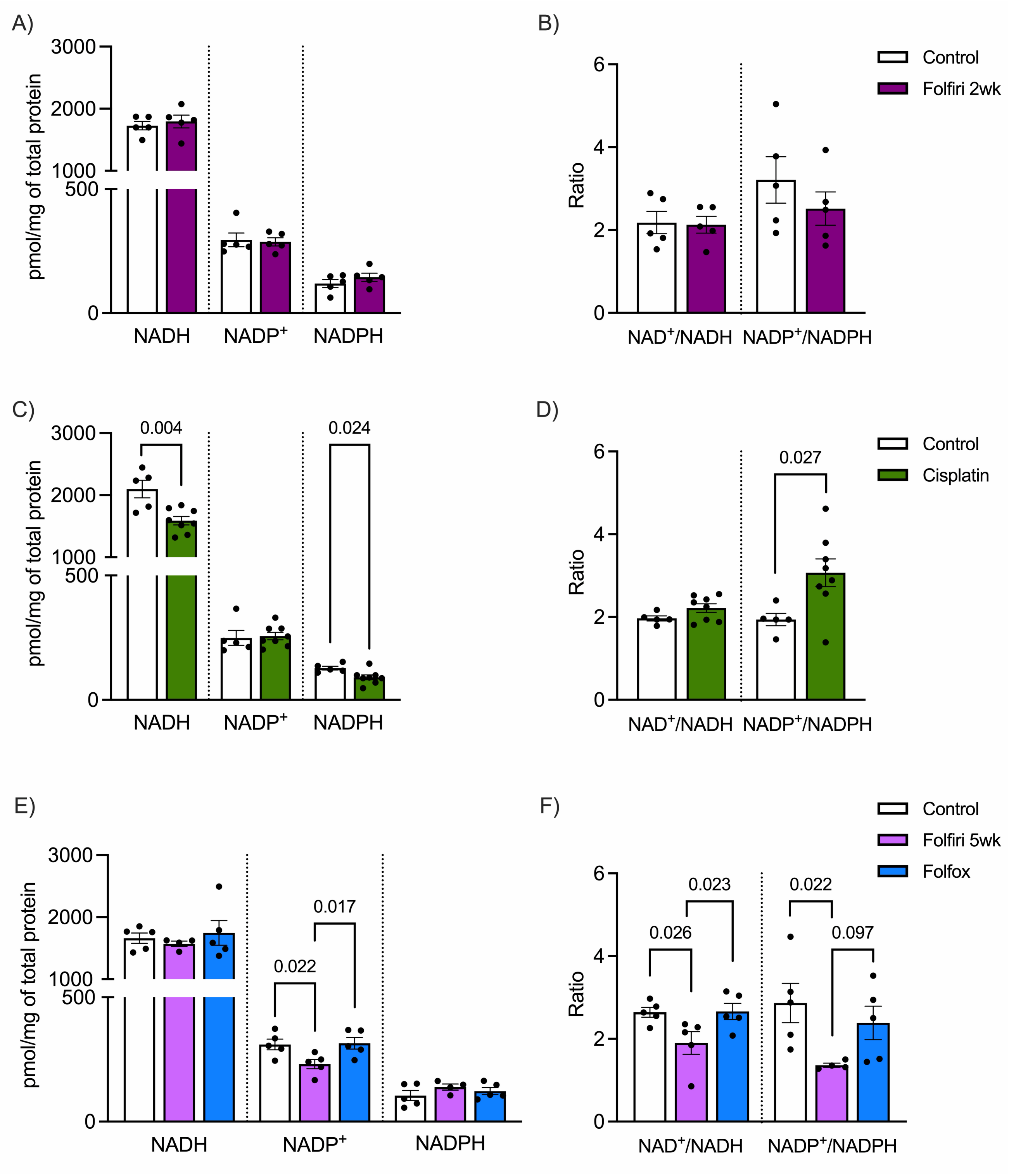


***Supplementary figure 2. Muscle NAD metabolites and metabolite ratios, related to Figure 1.***
*NADH, NADP$^+$, and NADPH concentrations in quadriceps muscle tissue of folfiri (N=5) (A), cisplatin (N=8) (C), and the 5-week folfiri/folfox (N=5 per group) (E) experiments with respective controls (N=5) in each experiment.*
*NAD$^+$/NADH and NADP$^+$/NADPH ratios in muscle tissue of folfiri (N=5) (B), cisplatin (N=5) (D), and the 5-week folfiri/folfox (N=5 per group) (F) experiments with respective controls (N=5) in each experiment.*

*Data are shown as individual values and mean ± SEM. Statistical analysis was performed using two-tailed Student's t test or one-way ANOVA followed by Fisher's LSD test for parametric data, and Mann-Whitney test or Kruskal-Wallis test followed by uncorrected Dunn's test for nonparametric data.*

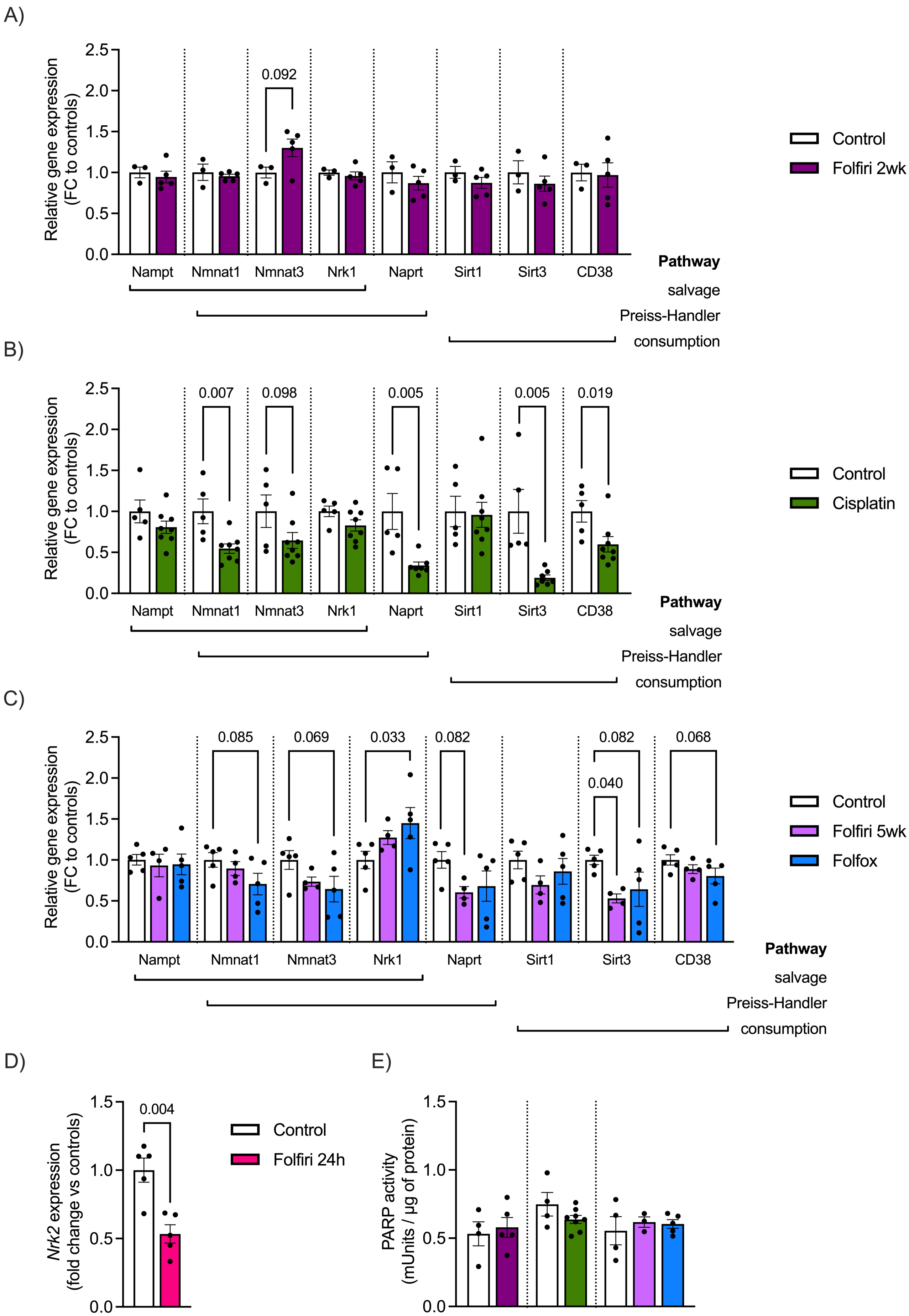

A)
Relative gene expression (FC to controls)
2.5
2.0
1.5
1.0
0.5
0.0
0.092
Nampt
Nmnat1
Nmnat3
Nrk1
Naprt
Sirt1
Sirt3
CD38
Pathway
salvage
Preiss-Handler
consumption
Control
Folfiri 2wk
B)
Relative gene expression (FC to controls)
0.007
0.098
0.005
0.005
0.019
Nampt
Nmnat1
Nmnat3
Nrk1
Naprt
Sirt1
Sirt3
CD38
Pathway
salvage
Preiss-Handler
consumption
Control
Cisplatin
C)
Relative gene expression (FC to controls)
0.085
0.069
0.033
0.082
0.082
0.040
0.068
Nampt
Nmnat1
Nmnat3
Nrk1
Naprt
Sirt1
Sirt3
CD38
Pathway
salvage
Preiss-Handler
consumption
Control
Folfiri 5wk
Folfox
D)
Nrk2 expression (fold change vs controls)
1.5
1.0
0.5
0.0
0.004
Control
Folfiri 24h
E)
PARP activity (mUnits / µg of protein)
1.5
1.0
0.5
0.0

***Supplementary figure 3. NAD$^+$ metabolism in the skeletal muscle of chemotherapy-administered healthy mice, related to Figure 1.***

*Relative mRNA expression of the NAD$^+$ biosynthetic and consuming enzymes in the quadriceps muscle of A) 2-week folfiri-treated healthy mice (N=5) expressed as fold change (FC) compared to controls (=1) (N=5), B) 2-week cisplatin-treated mice (N=8) and respective controls (N=5), and C) 5-week folfiri- (N=4) or folfox-treated mice (N=5) and respective controls (N=5).*

*D) Quadriceps muscle Nrk2 expression 24 hours post-folfiri administration (N=5).*

*E) Quadriceps muscle PARP activity in the 2- and 5-week chemotherapy experiments (N=3-8 per group).*

*Data are shown as individual values and mean ± SEM. Statistical analysis was performed using two-tailed Student's t test or one-way ANOVA followed by Fisher's LSD test for parametric data, and Mann-Whitney test or Kruskal-Wallis test followed by uncorrected Dunn's test for nonparametric data.*

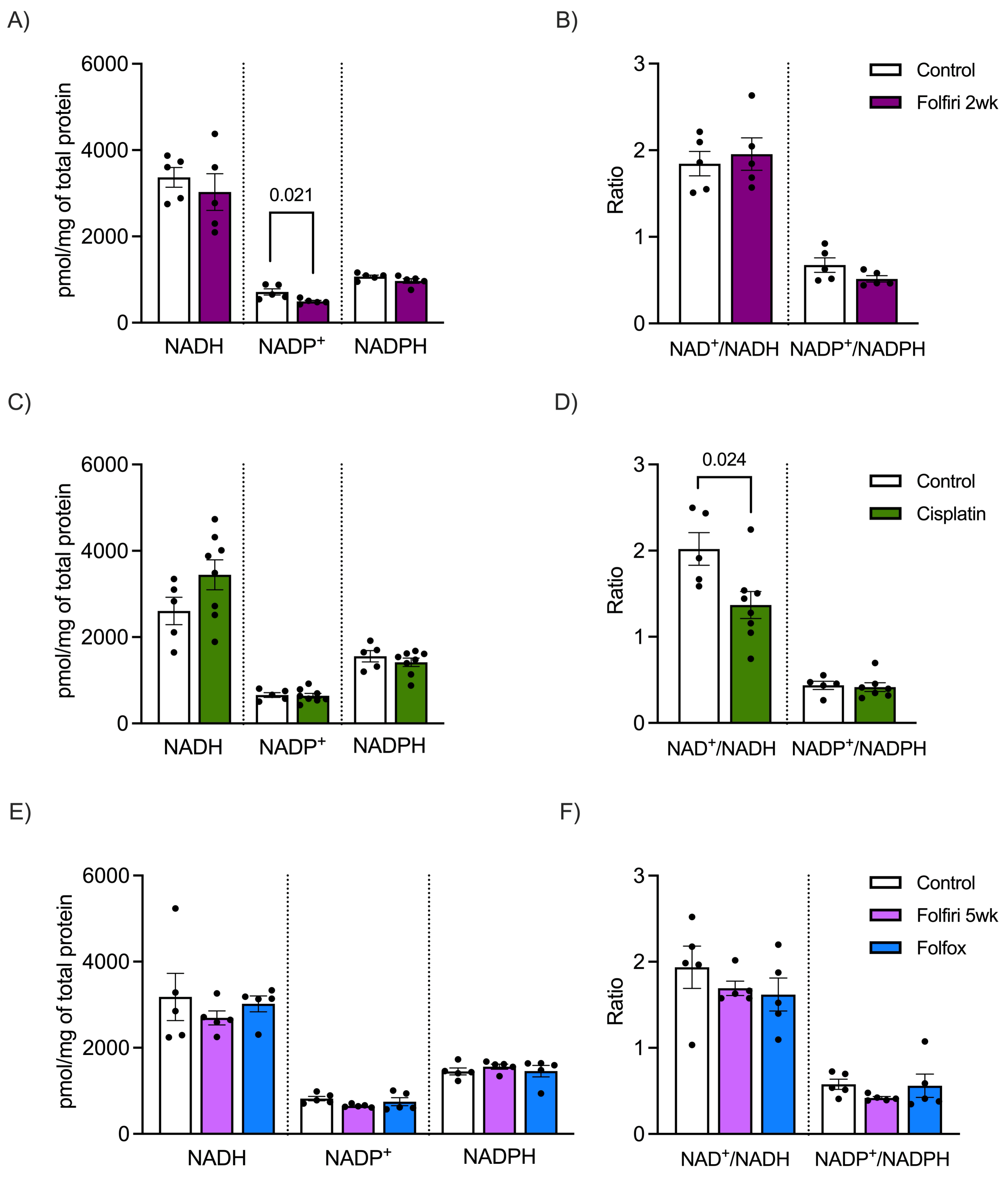


***Supplementary figure 4. Liver NAD metabolites and metabolite ratios, related to Figure 1.***
*NADH, NADP+, and NADPH concentrations in liver tissue of folfiri (N=5) (A), cisplatin (N=8) (C), and the 5-week folfiri/folfox (N=5 per group) (E) experiments with respective controls (N=5) in each experiment.*
*NAD+/NADH and NADP+/NADPH ratios in liver tissue of folfiri (N=5) (B), cisplatin (N=5) (D), and the 5-week folfiri/folfox (N=5 per group) (F) experiments with respective controls (N=5) in each experiment.*

*Data are shown as individual values and mean ± SEM. Statistical analysis was performed using two-tailed Student's t test or one-way ANOVA followed by Fisher's LSD test for parametric data, and Mann-Whitney test or Kruskal-Wallis test followed by uncorrected Dunn's test for nonparametric data.*

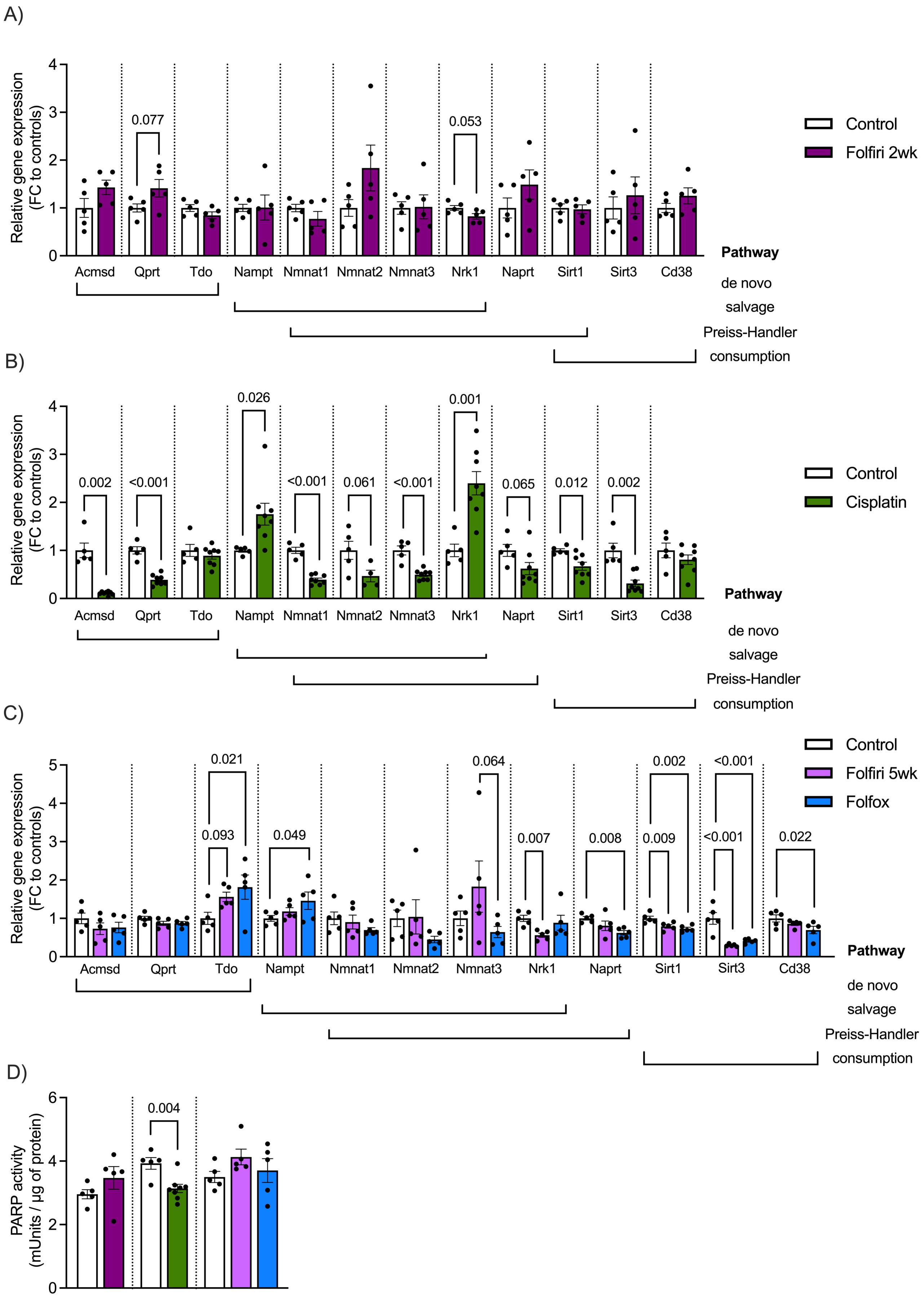

A)
Relative gene expression (FC to controls)
0.077
0.053
Control
Folfiri 2wk
Acmsd Qprt Tdo Nampt Nmnat1 Nmnat2 Nmnat3 Nrk1 Naprt Sirt1 Sirt3 Cd38
Pathway
de novo
salvage
Preiss-Handler
consumption
B)
Relative gene expression (FC to controls)
0.002 <0.001 0.026 <0.001 0.061 <0.001 0.001 0.065 0.012 0.002
Control
Cisplatin
Acmsd Qprt Tdo Nampt Nmnat1 Nmnat2 Nmnat3 Nrk1 Naprt Sirt1 Sirt3 Cd38
Pathway
de novo
salvage
Preiss-Handler
consumption
C)
Relative gene expression (FC to controls)
0.021 0.093 0.049 0.064 0.007 0.008 0.002 0.009 <0.001 <0.001 0.022
Control
Folfiri 5wk
Folfox
Acmsd Qprt Tdo Nampt Nmnat1 Nmnat2 Nmnat3 Nrk1 Naprt Sirt1 Sirt3 Cd38
Pathway
de novo
salvage
Preiss-Handler
consumption
D)
PARP activity (mUnits / µg of protein)
0.004

***Supplementary figure 5. NAD$^+$ metabolism in the liver of chemotherapy-administered healthy mice, related to Figure 1***.

*Relative mRNA expression of the NAD$^+$ biosynthetic and consuming enzymes in the liver of A) 2-week folfiri-treated healthy mice (N=5) expressed as fold change (FC) compared to controls (=1) (N=5), B) 2-week cisplatin-treated mice (N=8) and respective controls (N=5), and C) 5-week folfiri- (N=4) or folfox-treated mice (N=5) and respective controls (N=5).*

*D) Liver PARP activity in the 2- and 5-week chemotherapy experiments (N=5-8 per group).*

*Data are shown as individual values and mean ± SEM. Statistical analysis was performed using two-tailed Student's t test or one-way ANOVA followed by Fisher's LSD test for parametric data, and Mann-Whitney test or Kruskal-Wallis test followed by uncorrected Dunn's test for nonparametric data.*

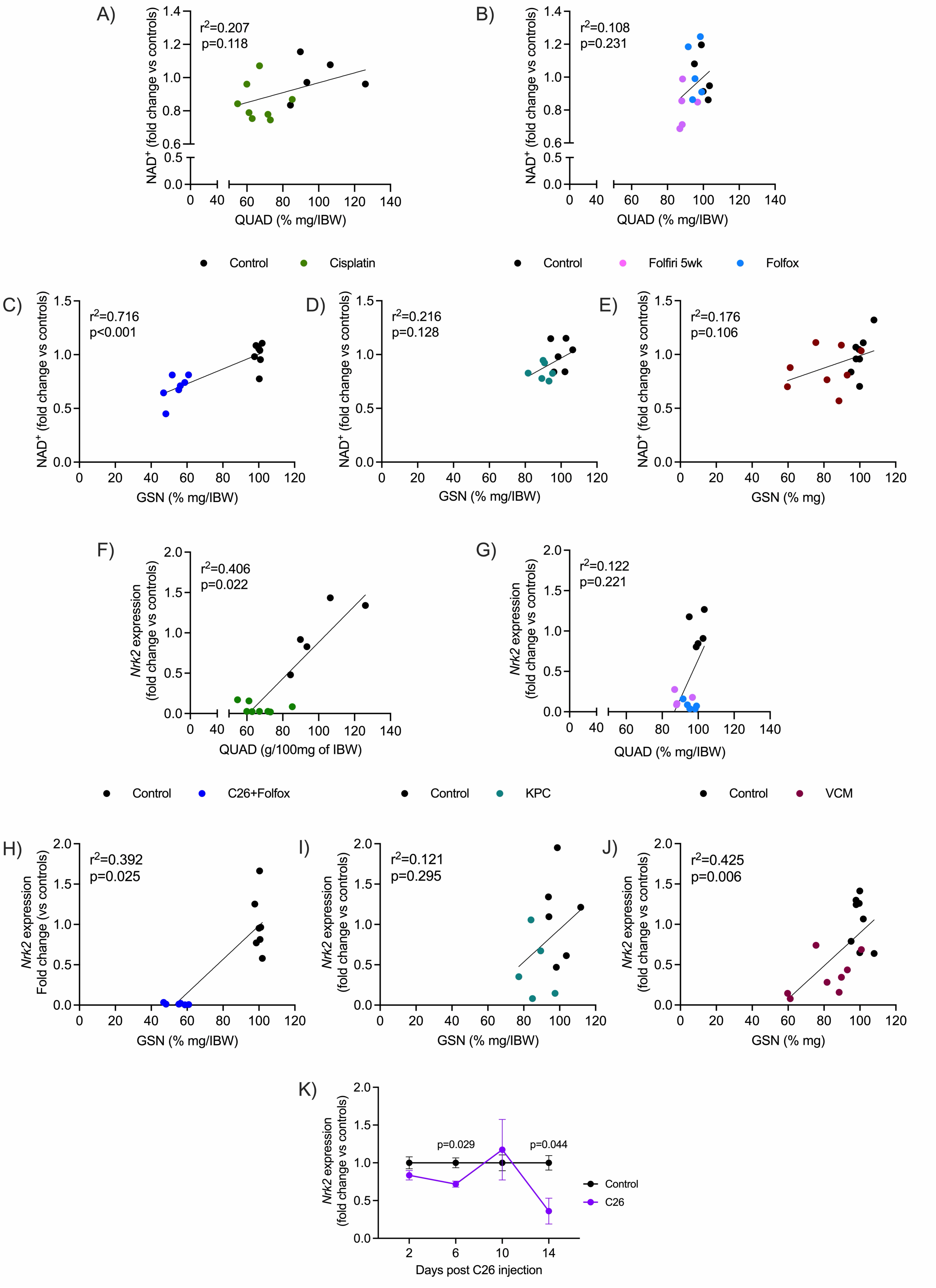
A)
r²=0.207
p=0.118
NAD⁺ (fold change vs controls)
QUAD (% mg/IBW)
B)
r²=0.108
p=0.231
NAD⁺ (fold change vs controls)
QUAD (% mg/IBW)
Control
Cisplatin
Control
Folfiri 5wk
Folfox
C)
r²=0.716
p<0.001
NAD⁺ (fold change vs controls)
GSN (% mg/IBW)
D)
r²=0.216
p=0.128
NAD⁺ (fold change vs controls)
GSN (% mg/IBW)
E)
r²=0.176
p=0.106
NAD⁺ (fold change vs controls)
GSN (% mg)
F)
r²=0.406
p=0.022
Nrk2 expression (fold change vs controls)
QUAD (g/100mg of IBW)
G)
r²=0.122
p=0.221
Nrk2 expression (fold change vs controls)
QUAD (% mg/IBW)
Control
C26+Folfox
Control
KPC
Control
VCM
H)
r²=0.392
p=0.025
Nrk2 expression Fold change (vs controls)
GSN (% mg/IBW)
I)
r²=0.121
p=0.295
Nrk2 expression (fold change vs controls)
GSN (% mg/IBW)
J)
r²=0.425
p=0.006
Nrk2 expression (fold change vs controls)
GSN (% mg)
K)
p=0.029
p=0.044
Nrk2 expression (fold change vs controls)
Days post C26 injection
Control
C26

***Supplementary figure 6. Correlations between NAD$^+$ or Nrk2 and muscle mass across individual experiments with muscle wasting: chemotherapy-induced cachexia and in previously published cancer- and cancer plus chemotherapy-induced cachexia (8). Related to Figure 2.*** *Correlations between NAD$^+$ and quadriceps mass (% mg per initial body weight (IBW)) are shown for A) the cisplatin experiment and B) the 5-week folfiri and folfox experiment.*

*Correlations between NAD$^+$ and gastrocnemius (GSN) muscle mass (% mg/IBW) are shown for C) C26-Folfox, D) KPC and E) VCM mouse models. Correlation between Nrk2 expression and quadriceps mass (% mg/IBW) in F) the cisplatin experiment and G) the 5-week folfiri and folfox experiment.*

*Correlation between Nrk2 and GSN muscle mass (% mg/IBW) are shown for H) C26-Folfox, I) KPC and J) VCM mouse models.*

*K) Timecourse of Nrk2 expression on days 2, 6, 10 and 14 in the skeletal muscle after C26 cell injection in mice.*

*Muscle mass is normalized by initial body weight (IBW) and represented as a percentage of the mean of the control group and NAD$^+$/Nrk2 levels are expressed as fold change versus controls. Statistical analyses included normality testing using the Shapiro-Willk test, followed by Pearson or Spearman correlation for parametric and nonparametric data, respectively. Welch's t test was used to analyze data from the C26 timecourse.*
